## Supplementary Information for "An adaptive biomolecular condensation response is conserved across environmentally divergent species"

#### Supplementary Figure S1.

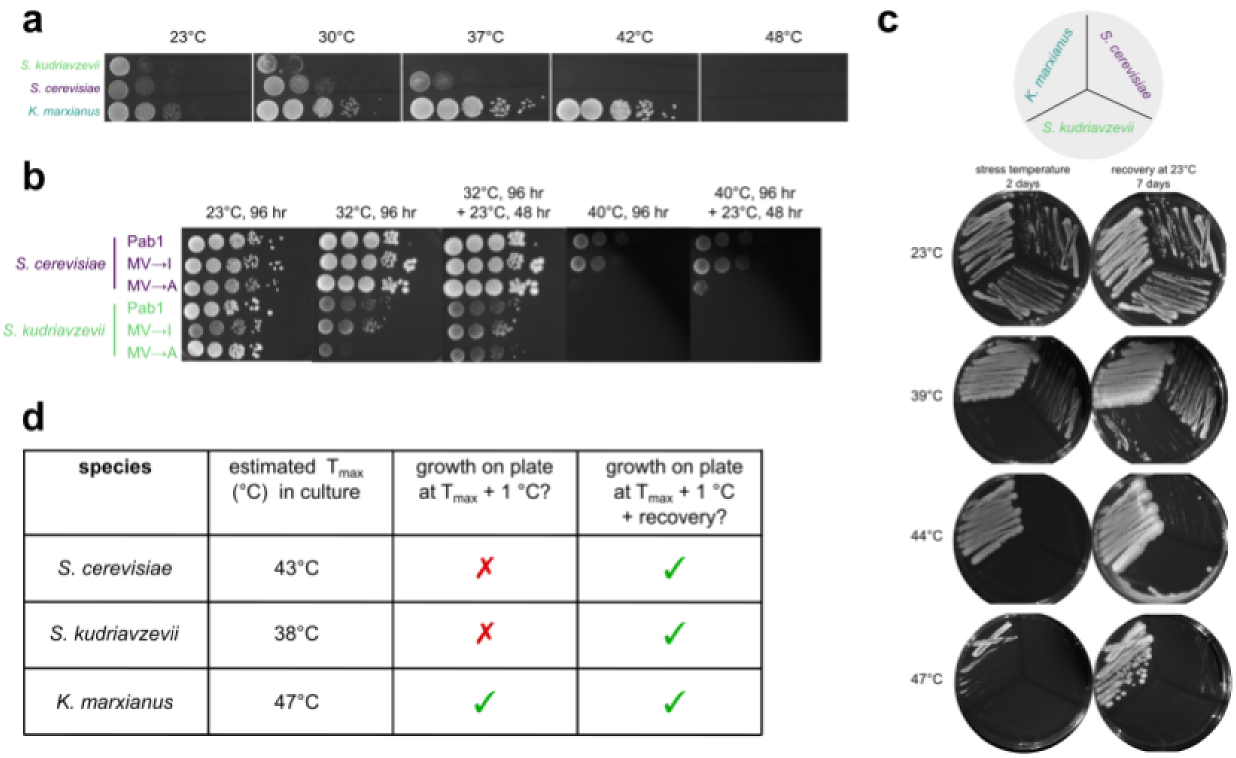


| species | estimated $T_{max}$ (°C) in culture | growth on plate at $T_{max}$ + 1 °C? | growth on plate at $T_{max}$ + 1 °C + recovery? |
|---|---|---|---|
| *S. cerevisiae* | 43°C | ✗ | ✓ |
| *S. kudriavzevii* | 38°C | ✗ | ✓ |
| *K. marxianus* | 47°C | ✓ | ✓ |

**Fig S1. Growth, recovery, and death phenotypes of each three species. a** Spot assays of S. cerevisiae, S. kudriavzevii, and K. marxianus strains. Plates were incubated at 23, 30, 37, 42, or 48°C for 2 days and then imaged. Columns are 10-fold dilutions. **b** Biological replicate of Figure 4e. **c** Growth on plates of each wild-type yeast species after a single-colony streak on a YPD plate. Each plate was incubated at either control or each species' $T_{max}$ +1°C (calculated from culture) for two days, imaged, then shifted to room temperature, grown for 7 days, and imaged again. **d** Summary table of panel c.

### Supplementary Figure S2.

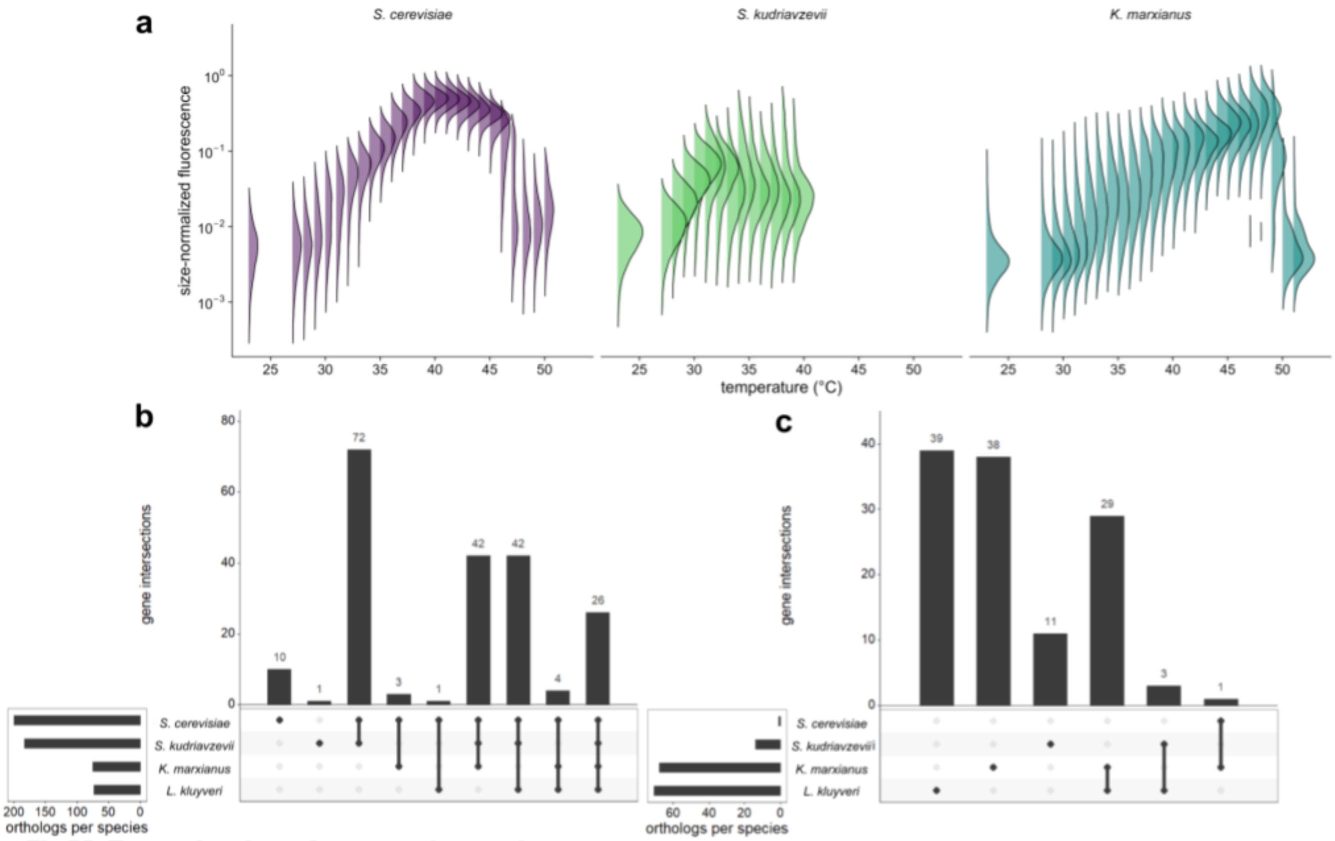


**Fig S2. Expression dynamics across temperatures.**
**a** Density distributions of red (endogenous SSA4-mCherry in *S. kudriavzevii* and *S. cerevisiae*) or green (plasmid-expressed SSA3p-eGFP in *K. marxianus*) fluorescence normalized by forward scatter for each species and temperature. **c**, **d** comparison of Msn2/4 up (**c**) and downregulated (**d**) genes from our study and *L. kluyveri* from Brion et al., 2016. There is strong overlap in upregulated genes and lack of overlap in downregulated genes between pre- and post-duplication relatives, consistent with the hypothesis that the post-duplication orthologs should be substantially similar if there are some ancestral genes which were only recruited into the Msn2/4 post-whole-genome duplication.

Supplementary Figure S3.

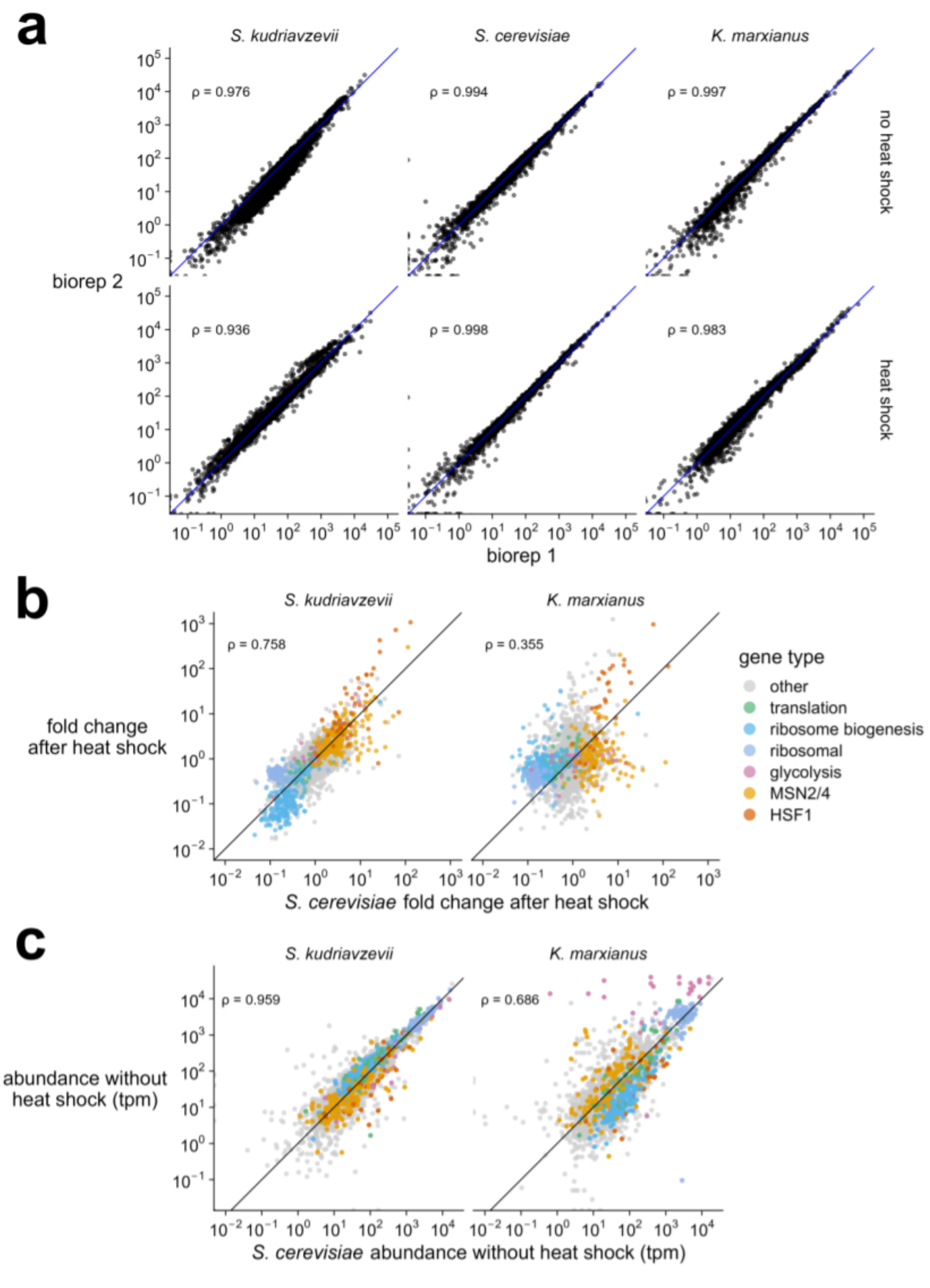


**Fig S3. Strong correlations in biological replicates and within treatment condition comparisons.** **a** Transcript abundance (transcripts per million, tpm) between biological replicates in each species. Correlations are represented by Pearson's rho (ρ) calculated between replicates. **b** Fold change distribution for groups of genes (colored by gene type) after stress in each species. Correlations are represented by Pearson's rho (ρ) calculated between species. **c** Transcript abundance (transcripts per million, tpm) in each species without a heat shock. Correlations are represented by Pearson's rho (ρ) calculated between species.

### Table 1.

| Well | Protein | baseline_radius | baseline_radius_sd | sd_check |
|---|---|---|---|---|
| F2 | SkPab1WT | 4.907570444444445 | 0.10385392527609397 | TRUE |
| H3 | KmPab1MV.I | 5.020872173913044 | 0.0895518637924171 | TRUE |
| K4 | KmPab1WT | 4.873762608695652 | 0.14987312516735884 | TRUE |
| K5 | KmPab1MV.A | 4.745623125 | 0.0936844469282388 | TRUE |
| K9 | SkPab1MV.A | 4.7453607692307695 | 0.09255458339646018 | TRUE |
| L12 | SkPab1MV.I | 5.165217826086956 | 0.232538851357174 | TRUE |
| NA | J-ScPab1MV.A | 4.314985806451613 | 0.07610316652445605 | TRUE |
| NA | J-ScPab1MV.I | 4.351886913580247 | 0.05017632628147393 | TRUE |
| NA | J-ScPab1WT | 4.392046875 | 0.08042537087253118 | TRUE |

**Table 1. Pab1 baseline size estimations.** Well indicates DLS experimental well; Protein indicates species abbreviation + Pab1 + mutant version;  baseline_radius shows value of the mean radius in nm of measurements below 35°C; baseline_radius_sd indicates the standard deviation of baseline_radius under 35°C; sd_check: TRUE if baseline_radius_sd is less than 5% of baseline_radius. Data for *S. cerevisiae* are from (Riback et al. 2017).

### Table 2.

| Well | Protein | $T_{demix}$ | Radius | Rep | Species | Mutant |
|---|---|---|---|---|---|---|
| F2 | SkPab1WT | 38.1963 | 10.4914 | rep_1 | Skud | WT |
| H3 | KmPab1MV.I | 46.0497 | 9.03576 | rep_1 | Kmarx | MV.I |
| K4 | KmPab1WT | 48.9093 | 10.2584 | rep_4 | Kmarx | WT |
| K5 | KmPab1MV.A | 50.8701 | 9.70903 | rep_2 | Kmarx | MV.A |
| K9 | SkPab1MV.A | 39.6627 | 11.0515 | rep_2 | Skud | MV.A |
| L12 | SkPab1MV.I | 36.5503 | 9.43427 | rep_4 | Skud | MV.I |
| NA | J-ScPab1MV.A | 40.7226 | 9.13283 | rep_1 | Scere | WT |
| NA | J-ScPab1MV.I | 39.2875 | 8.23502 | rep_1 | Scere | MV.I |
| NA | J-ScPab1WT | 42.7316 | 8.99349 | rep_1 | Scere | MV.A |

**Table 2. Pab1 $T_{condense}$ and estimations and size measurements.** Well indicates DLS experimental well; Protein indicates species abbreviation + Pab1 + mutant version; $T_{demix}$ represents the temperature where the radius is as close to double the average baseline value below 35°C; Radius is the measured diameter in nm of the radius and $T_{demix}$ temperature; rep indicates which replicate is used for the calculation; Species shows an abbreviation of species used; Mutant indicates the version of Pab1 mutant used. Data for *S. cerevisiae* are from (Riback et al. 2017).
